# Large-scale Deep Proteomic Analysis in Alzheimer’s Disease Brain Regions Across Race and Ethnicity

**DOI:** 10.1101/2024.04.22.590547

**Authors:** Fatemeh Seifar, Edward J. Fox, Anantharaman Shantaraman, Yue Liu, Eric B. Dammer, Erica Modeste, Duc M. Duong, Luming Yin, Adam N. Trautwig, Qi Guo, Kaiming Xu, Lingyan Ping, Joseph S. Reddy, Mariet Allen, Zachary Quicksall, Laura Heath, Jo Scanlan, Erming Wang, Minghui Wang, Abby Vander Linden, William Poehlman, Xianfeng Chen, Saurabh Baheti, Charlotte Ho, Thuy Nguyen, Geovanna Yepez, Adriana O. Mitchell, Stephanie R. Oatman, Xue Wang, Minerva M. Carrasquillo, Alexi Runnels, Thomas Beach, Geidy E. Serrano, Dennis W. Dickson, Edward B. Lee, Todd E. Golde, Stefan Prokop, Lisa L. Barnes, Bin Zhang, Varham Haroutunian, Marla Gearing, James J. Lah, Philip De Jager, David A Bennett, Anna Greenwood, Nilüfer Ertekin-Taner, Allan I. Levey, Aliza Wingo, Thomas Wingo, Nicholas T. Seyfried

## Abstract

**Introduction:** Alzheimer’s disease (AD) is the most prevalent neurodegenerative disease, yet our comprehension predominantly relies on studies within the non-Hispanic White (NHW) population. Here we aimed to provide comprehensive insights into the proteomic landscape of AD across diverse racial and ethnic groups.

**Methods:** Dorsolateral prefrontal cortex (DLPFC) and superior temporal gyrus (STG) brain tissues were donated from multiple centers (Mayo Clinic, Emory University, Rush University, Mt. Sinai School of Medicine) and were harmonized through neuropathological evaluation, specifically adhering to the Braak staging and CERAD criteria. Among 1105 DLPFC tissue samples (998 unique individuals), 333 were from African American donors, 223 from Latino Americans, 529 from NHW donors, and the rest were from a mixed or unknown racial background. Among 280 STG tissue samples (244 unique individuals), 86 were African American, 76 Latino American, 116 NHW and the rest were mixed or unknown ethnicity. All tissues were uniformly homogenized and analyzed by tandem mass tag mass spectrometry (TMT-MS).

**Results:** As a Quality control (QC) measure, proteins with more than 50% missing values were removed and iterative principal component analysis was conducted to remove outliers within brain regions. After QC, 9,180 and 9,734 proteins remained in the DLPC and STG proteome, respectively, of which approximately 9,000 proteins were shared between regions. Protein levels of microtubule-associated protein tau (MAPT) and amyloid-precursor protein (APP) demonstrated AD-related elevations in DLPFC tissues with a strong association with CERAD and Braak across racial groups. APOE4 protein levels in brain were highly concordant with *APOE* genotype of the individuals.

**Discussion:** This comprehensive region resolved large-scale proteomic dataset provides a resource for the understanding of ethnoracial-specific protein differences in AD brain.

## 1. Background

Alzheimer’s disease (AD) presents a significant global health challenge, with its prevalence affecting millions worldwide (1, 2). Notably, African Americans and Hispanic Americans are almost twice as likely to have AD and other dementias compared to Caucasians (3, 4). The mechanisms contributing to this disparity may be multifaceted, involving a combination of genetic differences across race and ethnicity, as well as societal and environmental inequities that disproportionately affect African Americans and Latino Americans (4–6). These include lower levels and quality of education, as well as higher rates of poverty (7–12). To date, the bulk of our knowledge regarding AD pathophysiology derives from studies conducted within the non-Hispanic White (NHW) population. Emerging evidence suggests differences in some molecular measures, such as lower cerebrospinal fluid (CSF) levels of Tau and other synaptic proteins in African Americans with AD compared to NHWs (13–16). However, the extent of similarities or differences for AD-related molecular perturbations in postmortem brain tissues from diverse populations remain incompletely understood. Consequently, there is a significant gap in our understanding of the ethnoracial disparities inherent in the pathophysiology of AD. To address this gap in knowledge, the National Institute on Aging and Accelerating Medicines Partnership in AD (AMP-AD) aimed to promote inclusivity in multi-omics AD research, to unravel unique molecular signatures and pathways (17). This step is crucial for achieving a more precise definition of AD that accounts for variations across racial, ethnic, and genetic backgrounds.

Proteins serve as optimal markers for understanding “proteinopathies” like AD and other neurodegenerative disease due to their proximity to pathologic and phenotypic changes in disease (18). With the advancement of multiplex isobaric tandem mass tags (TMT), off-line fractionation, and high-resolution mass spectrometry, proteomic datasets are now approaching the depth of transcriptomic datasets (19–21). Integrated analyses of proteomic and transcriptomic data in AD post-mortem brain cohorts indicate that these approaches can yield both complementary and unique insights in human brain (22–24). However, a comprehensive and detailed proteome atlas of the human brain spanning various regions, races, and ethnicities is still lacking. Such studies could uncover race-specific protein differences, shedding light on distinct pathways, pathophysiologies, biomarkers, and potential therapeutic targets in AD.

Using TMT coupled with mass spectrometry (TMT-MS) we report a large-scale and deep proteome (∼10,000 proteins) of the post-mortem dorsolateral prefrontal cortex (DLPFC) from 998 individuals and the superior temporal gyrus (STG) from 244 individuals across control and pathologically defined AD cases. Of these, approximately 50% of the samples were from racially and ethnically diverse brain donors. Implementing a methodology for quality control measures, we ensured the removal of batch-related variations from the dataset. Subsequently, variance partitioning analyses were carried out to identify top proteins based on individual characteristics, such as sex, race, and AD diagnosis, across both DLPFC and STG tissues. Through TMT-MS, we characterized core proteins associated with AD pathology, including amyloid precursor protein (APP) and the microtubule-associated protein tau (MAPT), revealing a clear correlation of APP and MAPT levels with CERAD and Braak stages. Furthermore, we show consistency between APOE4 protein levels with *APOE4* genotyping in brain. This comprehensive large-scale proteomic dataset not only demonstrates the validity of our findings but also establishes the foundation for a better understanding of ethnoracial-specific protein modulations, distinct pathways, pathologies, biomarkers, and potential therapeutic targets in AD.

## 2. Methods

### 2.1. Brain Tissue Collection and Cohort Characteristics

The proteomics data utilized in this study were a part of AMP-AD Diversity Initiative, a collaborative effort involving multiple research sites. The comprehensive dataset includes information from different multi-omics data including proteomics, genomics, and metabolomics. While the mass spectrometry and case selection analysis has been extensively described in the data descriptor manuscript (25), this study specifically focuses on database search, quality control (QC) and technical validation of the proteomics data.

In brief, brain samples were collected with the involvement of four institutions or data contribution sites: Mayo Clinic, Rush University, Mount Sinai University Hospital and Emory University for proteomics studies. The goal of this initiative is to include diverse contributions from African American and Latino American populations. Each of the data contribution sites gathered brain samples from affiliated brain banks, cohort studies, and AD Research Centers (ADRC) and were sent to Emory proteomics core for proteomic processing. A total of 1105 DLPFC tissues from 998 individuals were sent from all four data contribution sites including n= 129 from Emory University (including 22 samples from University of Pennsylvania), n= 399 from Mayo Clinic, n= 205 from Mount Sinai University Hospital and n= 372 from Rush University. Frontal brain tissues from each contribution site were processed separately from the others. In addition, amongst Emory samples, 26 samples from Mount Sinai University were replicated and among Mayo Clinic samples, 81 were replicated from Emory University samples.

A total of 280 superior temporal gyrus (STG) tissues from 244 individuals were obtained from Emory University (n= 129) and Mayo Clinic (n= 151), and both were processed simultaneously. Tissue homogenization, protein digestion, TMT peptide labeling, pH fractionation and liquid chromatography (LC)-MS/ MS are described in detail in the previous publication (25).

Collectively, LC-MS/MS led to a total of 6479 raw files from frontal cortex, and 1824 raw files from temporal cortex tissue samples (**Fig. 1A**), with the distribution as follows: Emory University Frontal Cortex Cohort: 431; Mayo Clinic Frontal Cortex Cohort: 2304; Mount Sinai Frontal Cortex Cohort: 1344; Rush University Frontal Cortex Cohort: 2400; and Emory University and Mayo Clinic Temporal Cortex Cohort: 1824.

**Fig 1.**
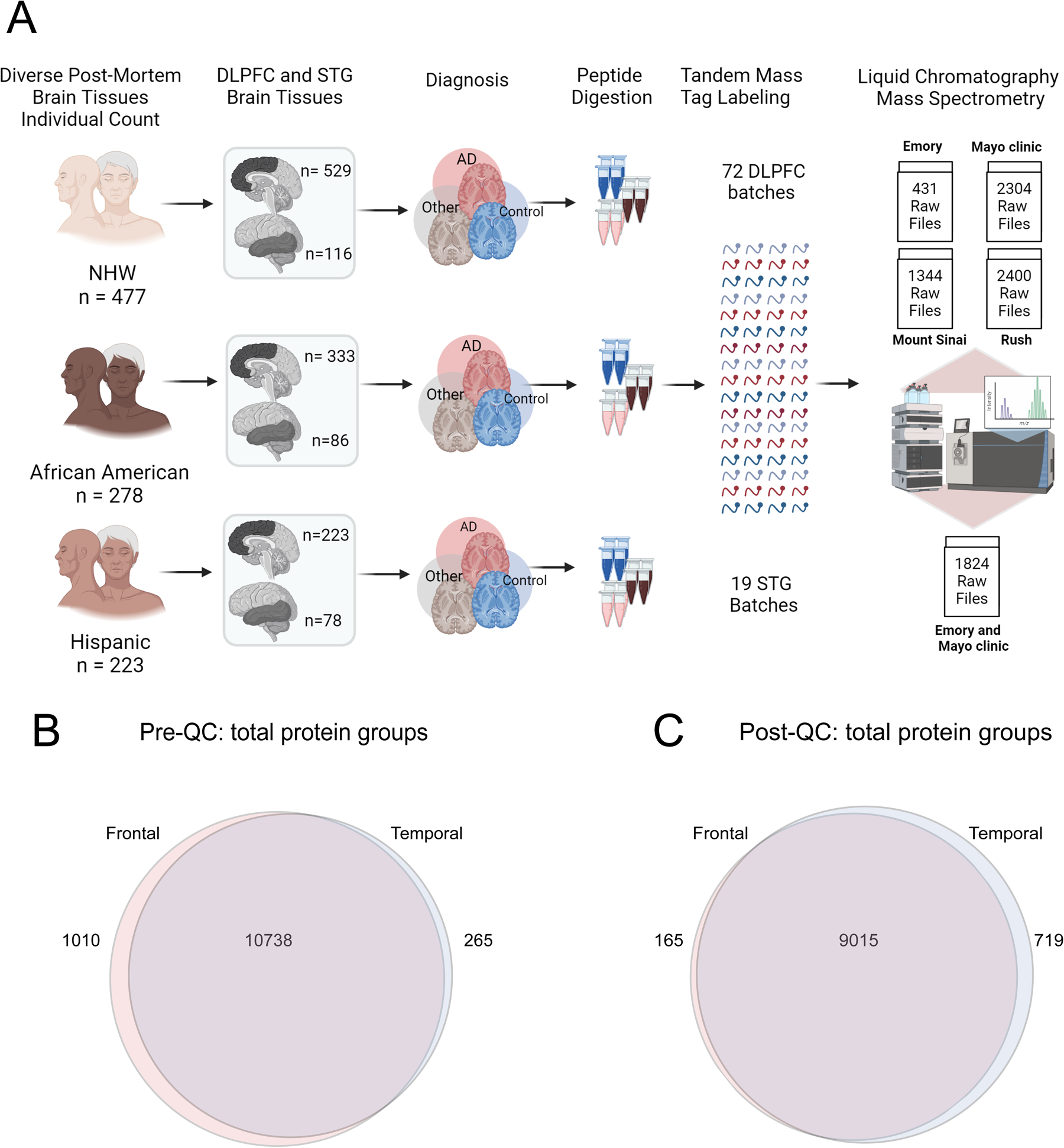
A. Schematic illustrating the cohort characteristics and the experimental workflow for mass spectrometry (MS) of the human brain proteome across frontal and temporal brain tissue samples. This study incorporated a total of 1105 dorsolateral prefrontal cortex (DLPFC) brain tissues from 998 individuals, categorized as follows: 529 non-Hispanic white (NHW), 333 African American, 223 Latino American, and others (n= 20) as applicable. These samples were sourced from four prominent data distribution sites: Emory University, Mayo Clinic, Rush University, and Mount Sinai University Hospital. Additionally, 280 STG tissues from a subset of 244 individuals were included, with 116 NHW, 86 African American, 78 Hispanic, and others as applicable. STG samples were obtained from a racially diverse set of specimens originating from Mayo Clinic and Emory, distributed across 19 batches. Tissues underwent an experimental pipeline involving protein digestion, batch randomization, TMT labeling, fractionation, and subsequent mass spectrometric measurements. A total of 72 DLPFC batches were processed, comprising 9 batches from Emory, 24 from Mayo Clinic, 14 from Mount Sinai, and 25 from Rush (comprising a total of 72 batches). The randomization of batches was conducted to ensure a representative and diverse dataset. The output included a total of 6479 raw files for DLPFC samples and 1824 raw files for STG. B. Venn diagram of total number of proteins quantified from DLPFC and STG samples. A total of 11748 protein groups were identified from DLPFC and 11003 from STG samples, with 10738 shared protein groups. C. Venn diagram of total protein from DLPFC and STG samples after quality control (QC) across all samples. 9180 protein groups were identified from DLPFC samples and 9734 from STG, with 9015 shared protein groups.

### 2.2. Database Searches and Protein Quantification

All raw files underwent a database search using Fragpipe (version 19.0) for DLPFC and STG datasets, separately. The database search parameters have been described (26, 27). Initially, mzML files were created from the original MS .raw files for frontal (6479 raw files across 72 batches) and temporal (1824 raw files across 19 batches) using the ProteoWizard MSConvert tool (version 3.0) with specific options, including ‘Write index,’ ‘TPP compatibility,’ ‘Use zlib compression,’ and a “peakPicking” filter setting.

Following the creation of mzML files for each set, they were subjected to a search using MSFragger (version 3.5). The human proteome database used contained 20,402 sequences (Swiss-Prot, downloaded 2/11/2019) along with corresponding decoys and common contaminants. The sequences include additional specific peptide sequences for the APOE ε4 and APOE ε2 alleles (28).

The search settings included a precursor mass tolerance of -20 to 20 ppm, a fragment mass tolerance of 20 ppm, mass calibration, parameter optimization, isotope error set to -1/0/1/2/3, strict-trypsin enzyme specificity, and allowance for up to two missed cleavages. Fully enzymatic cleavage type, peptide length (7 to 50), and peptide mass (200 to 5,000 Da) criteria were defined. Variable modifications included oxidation on methionine, N-terminal acetylation on protein, and TMTpro modification on the peptide N-terminus, with a maximum of 3 variable modifications per peptide. Static modifications comprised isobaric TMTpro (TMT16) modifications on lysine, along with carbamidomethylation of cysteine.

Post-MSFragger (version 3.6) search, Percolator (29) was used for PSM validation, succeeded by Philosopher (version 4.6.0) for protein inference using ProteinProphet and FDR filtering. Reports containing quantified peptides and UniprotID-identified proteins with an FDR < 1% were generated. The database search culminated in the identification of a total of 11,748 protein groups from frontal cortex samples and 11,003 from temporal cortex samples, revealing a shared set of 10,738 protein groups (**Fig. 1B**).

### 2.3. Data Analysis and QC

The data analysis, using R statistical software (version 4.3.2), adhered to a three-step process to ensure the consistency of the frontal (DLPC) and temporal (STG) tissue proteomic datasets.

The analysis workflow for data QC is illustrated in the flowcharts of **Fig 2A** and **Fig. 2B** in 3 main steps:

**Fig 2.**
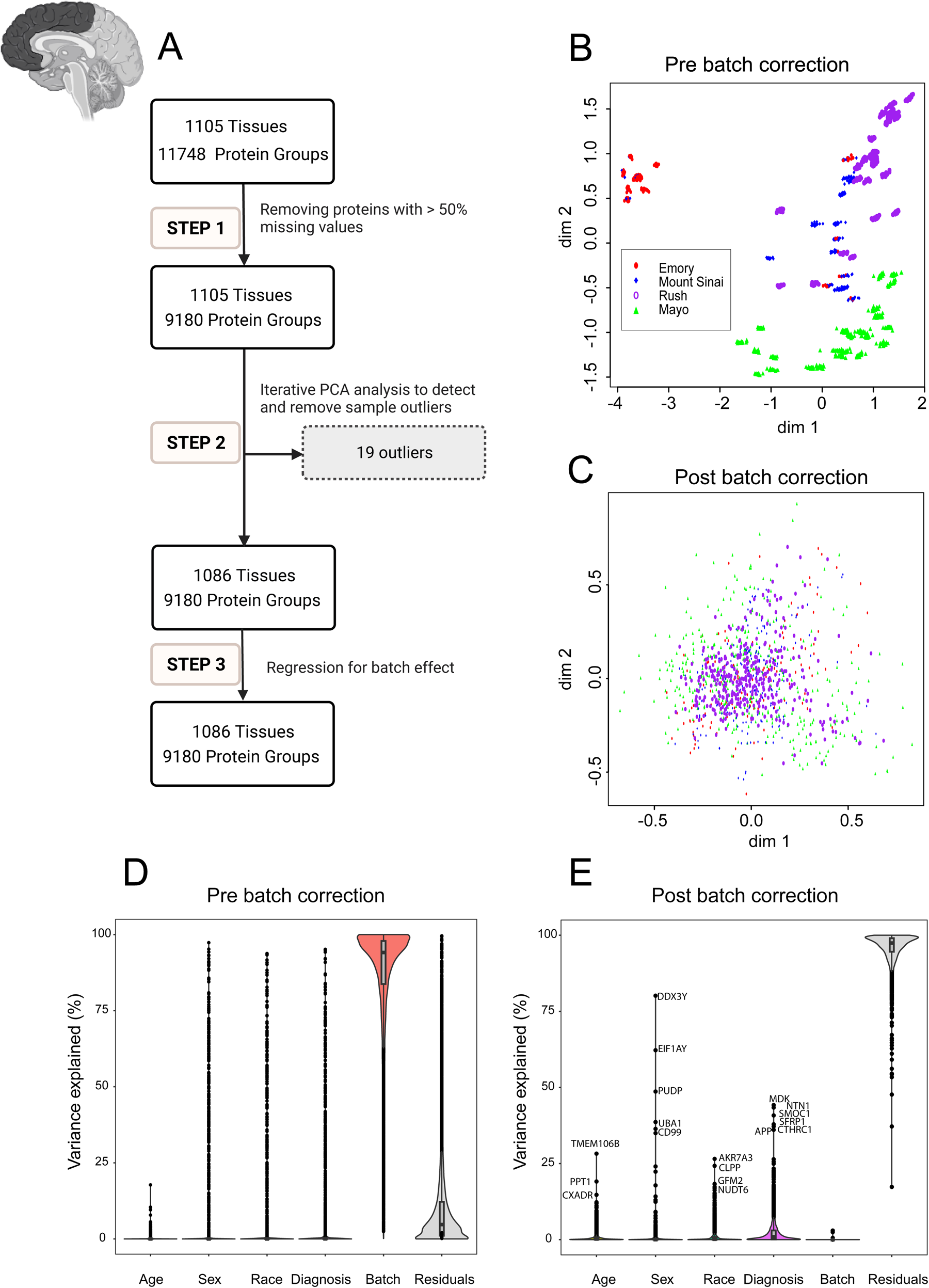
Quality Control (QC) and Batch Correction for DLPFC Tissue proteins. **A.** The QC workflow is illustrated in the flowchart in 3 main steps: Step 1. Pre-processing for missing values: Only proteins with missing data in less than 50% of the samples were retained. The ratio of protein abundance to the total protein abundance for each sample was calculated to adjust for sample loading differences resulting in 9180 proteins being retained across 1105 samples. Subsequently, the data was log2 transformation Step 2. Outlier detection and removal: Iterative principal component analysis (PCA) was employed to identify and eliminate sample outliers. After multiple rounds of PCA analysis, 19 outliers were identified and removed, leaving 9180 proteins across 1086 samples. Step 3. Batch effect regression: Variance attributable to batching was mitigated through regression of the 9180 proteins in 1086 samples. **B and C.** Multidimensional scaling (MDS) plot showing variation among samples (**B**) before correcting for batch and (**C**) after regressing for batch effect. The plot dimensions (dim 1 and 2) reveal distinctive clusters formed by samples by site (Emory (red), Mount Sinai (blue), Rush (purple), and Mayo (green)), with some scattering observed among samples before regressing for batch effect (**B**). (**C**) The plot illustrates the successful removal of variance due to batch. After correcting for batch effects, samples from all four sites - Emory (red), Mount Sinai (blue), Rush (purple), and Mayo (green) – cluster together, indicating a more cohesive grouping (n.b the change in scale from B to C). The correction mitigates the dispersion observed in panel **B**, highlighting the effectiveness of the batch correction procedure in harmonizing the sample distribution across different data distribution sites. **D and E.** Variance partition analysis using experimental factors to evaluate the percentage of explained variance in proteomic samples. Violin plots before (**D**) and after (**E**) batch correction illustrate the distribution of explained variances in overall proteomic values. The Y-axis represents the percentage of explained variance, while the X-axis depicts factors contributing to variance, such as age, sex, race, diagnosis, residuals, and batch. Notably, batch variance is present before batch correction, influencing the overall proteomic profile. Panel **E** displays the same factors on the X-axis after batch correction. Significantly, the violin plot demonstrates a substantial reduction in variance associated with batch, ultimately reaching near zero percent after batch regression. Moreover, even after batch correction, factors such as age, sex, race, AD diagnosis, and other individual traits (residual) had levels of impact on protein abundance patterns. Each point on the violin plot represents a specific protein, with the corresponding name next to it. This underscores the efficacy of the correction procedure in eliminating batch-related variability from the proteomic data.

#### 2.3.1. Step 1. Pre-processing for Missing Values

Proteins with missing data in less than 50% of the samples were retained as described (22, 30). The ratio of protein abundance to the total protein abundance for each sample was calculated to adjust for sample loading differences. Subsequently, a log_2_ transformation was applied to enhance the normality of the distribution of protein abundance, addressing potential skewness and stabilizing variance across samples.

#### 2.3.2. Step 2. Outlier Detection and Removal

Iterative principal component analysis (PCA) was employed to identify and eliminate samples more than 4 standard deviations from the mean of either the first or second principal component as previously described (31, 32). Multiple iterations of PCA were conducted, with outliers from each round being systematically removed before initiating the subsequent iteration. For the DLPFC, 19 outliers were removed after 5 rounds of PCA analysis. For the STG, two outliers were removed after three rounds of PCA analysis.

#### 2.3.3. Step 3. Accounting for Batch Effect

Briefly, a linear regression model was fitted to estimate the effect of protein sequencing batch. We then regressed out the batch effect from the protein abundance before the next step of analysis to minimize batch effects and enhance the reliability of downstream analyses. Following the QC process, a total of 9,180 proteins remained for the DLPFC and 9,734 proteins for the STG, with an overlap of 9,015 protein groups shared between the two brain regions (**Fig. 1C**).

## 3. Results

### 3.1. Proteomics Data Quality Control in Frontal and Temporal cortices

In large-scale TMT-MS proteomics studies, batch effects are inevitable due to technical reasons, especially when processing large cohorts in multiple separate batches (33, 34). To investigate variability and clustering patterns associated with batch effects among frontal cortex samples prior to normalization and batch correction, we employed Multidimensional scaling (MDS) plot. MDS is similar to PCA, which is used for visualizing high-dimensional data in lower-dimensional spaces (35). Before batch regression, distinctive clusters by samples from different sites were observed (**Fig. 2B**). However, after batch correction, samples clustered together, indicating the success of batch regression giving even distribution of data without regard to data distribution sites (**Fig. 2C**). The effectiveness of batch correction was also assessed through variance partition analysis, which revealed that the percentage of variance in protein abundance explained by batch was nearly zero after batch regression (**Fig. 2D**, **Fig. 2E**).

Similarly, to investigate the impact of batch on temporal cortex samples, MDS plots were utilized on post-search and post QC proteomics data. The plots illustrated a distinct clustering before QC, followed by an even distribution post-QC, underscoring the effectiveness of batch correction (**Fig. 3B**, **Fig. 3C**). In addition, batch variance revealed a high impact on the proteomic profile before correction in variance partitioning (**Fig. 3D**) and a substantial reduction in variance associated with batch after QC (**Fig. 3E**).

**Fig 3.**
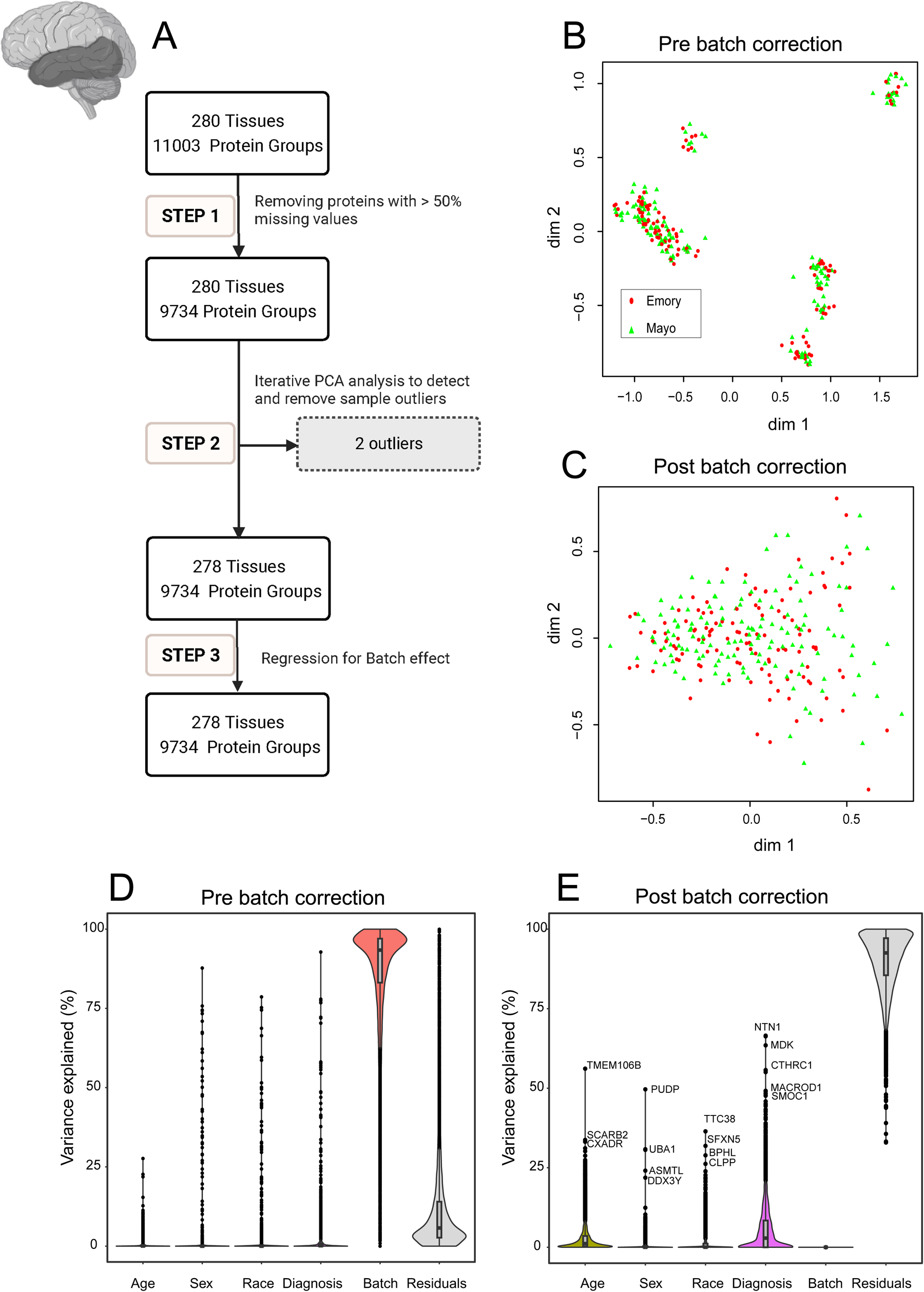
Quality Control (QC) and Batch Correction for STG Tissue proteins. **A.** The analysis workflow for data QC is depicted in three main steps: Step 1. Handling missing values: Proteins with missing data in more than 50% of the samples were removed, adjusting for sample loading differences through ratio calculation and log2 transformation. This yielded 9,734 proteins across 280 samples. Step 2. Identification and removal of outliers: Iterative principal component analysis (PCA) was utilized to detect and eliminate sample outliers. Following three rounds of PCA, two outliers were removed, resulting in 9,734 proteins across 278 samples. Step 3. Batch effect removal: Regression was applied to mitigate batch effects for the 9,734 proteins in 278 samples. **B and C.** Analysis of Multidimensional Scaling (MDS) plots: MDS plots depict sample variation (**B**) before batch correction and (**C**) after regression for batch effect. Emory (red) and Mayo (green) samples form distinctive clusters, with some scattering observed among samples before batch regression (**B**). (**C**) demonstrates the impact of batch regression, revealing a more cohesive grouping of Emory (red) and Mayo (green) samples. The correction effectively reduces the dispersion observed in panel **B**. **D and E.** Variance partition analysis for proteomic samples: Violin plots (**D**) before and (**E**) after batch correction show the distribution of explained variances in overall proteomic values. Panel D’s Y-axis represents the percentage of explained variance, while the X-axis includes factors like age, sex, race, diagnosis, residuals, and batch. Similar to Fig 2**.D**, batch variance revealed a high impact on the proteomic profile before correction. Panel **E** displays the same factors after batch correction, demonstrating a substantial reduction in variance associated with batch. In addition, after batch correction, age, sex, race, AD diagnosis, and other individual characteristics (residuals) remain influential factors shaping protein abundance patterns. Each data point represents a unique protein, with the corresponding protein names provided adjacent to the top points. This highlights the success of the regression analysis in eliminating batch-related variability from the proteomic data.

### 3.2. Variance of Protein Abundance in Frontal and Temporal Cortex Explained by Individual Characteristics

To understand how sex, race, and AD diagnosis is associated with protein abundance in the DLPFC and STG samples, we also employed variance partition analysis. This approach aimed to uncover the molecular basis of sex-, race-, and AD-associated variations in the brain (**Fig. 4**, **Fig. 5**). In current study, the analysis was performed using the traits available on synapse. Additional traits for Rush samples were provided separately by Rush Alzheimer’s Disease Research Center (https://www.radc.rush.edu/).

**Fig 4.**
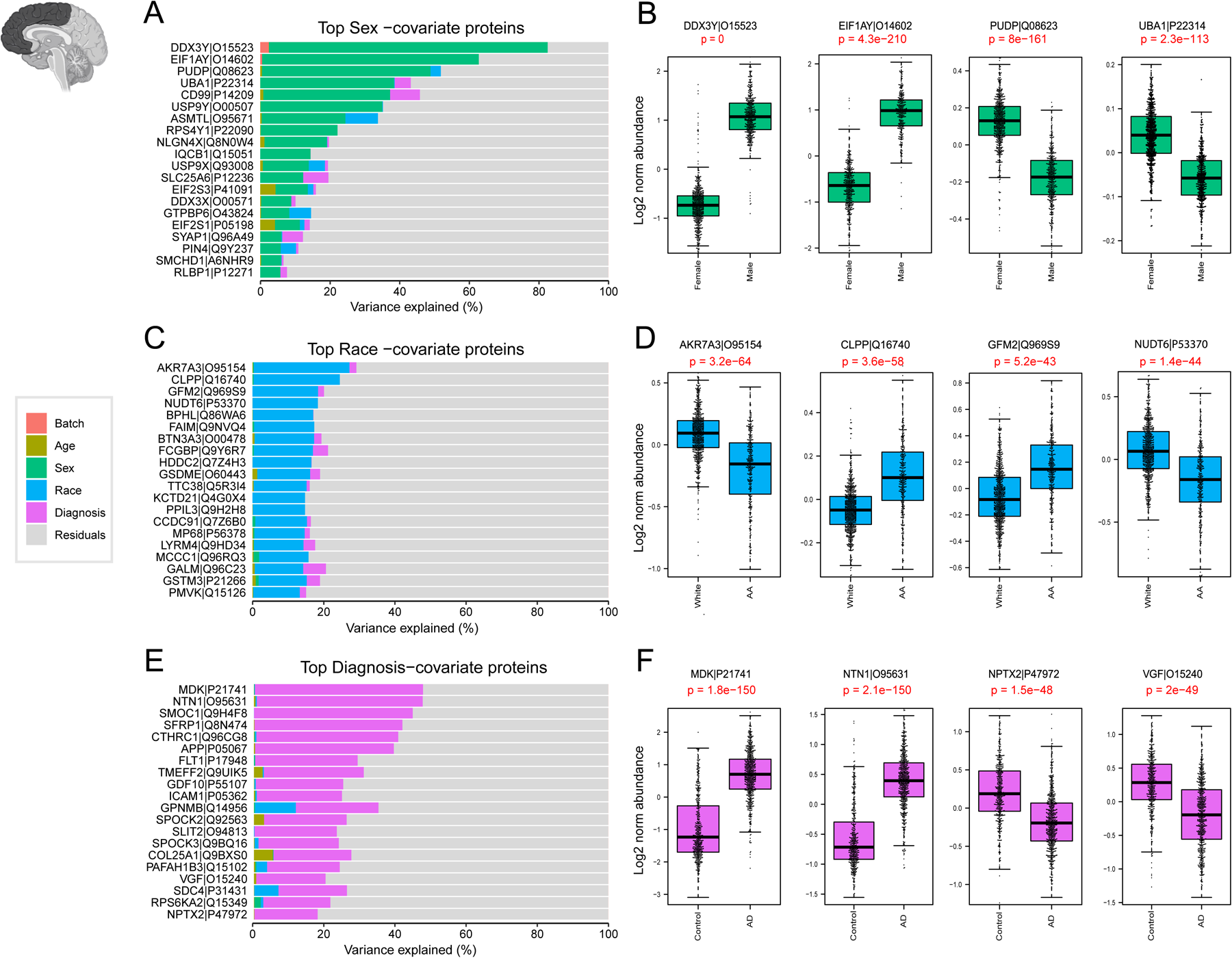
Variance Explained by individual Characteristics in DLPFC Tissues. The bar plots (**A, C, E**) depict the amount of variance explained by sex, race, and Alzheimer’s disease (AD) diagnosis across all DLPFC samples. **A.** Top-ranking proteins associated with sex in the dataset were identified through variance partitioning and depicted as bar plots. Boxplots in panel **B** illustrate the log2 normal abundance levels of four selected proteins exhibiting significant differences between males and females. These proteins serve as key indicators of sex-related variations and are depicted with statistical significance (p <0.05). **C.** Bar plots of top-ranking proteins associated with race differences in the DLPFC dataset. Boxplots in panel **D** illustrates the log2 normal abundance levels of four selected proteins demonstrating significant differences between African American individuals and other races (p <0.05). **E.** Bar plots identified top-ranking proteins contributing to the differences in the diagnosis of AD within the dataset. Boxplots in panel **F** display the log2 normal abundance levels of four selected proteins exhibiting significant differences between AD patients and controls, as well as other diagnostic categories (p <0.05).

**Fig 5.**
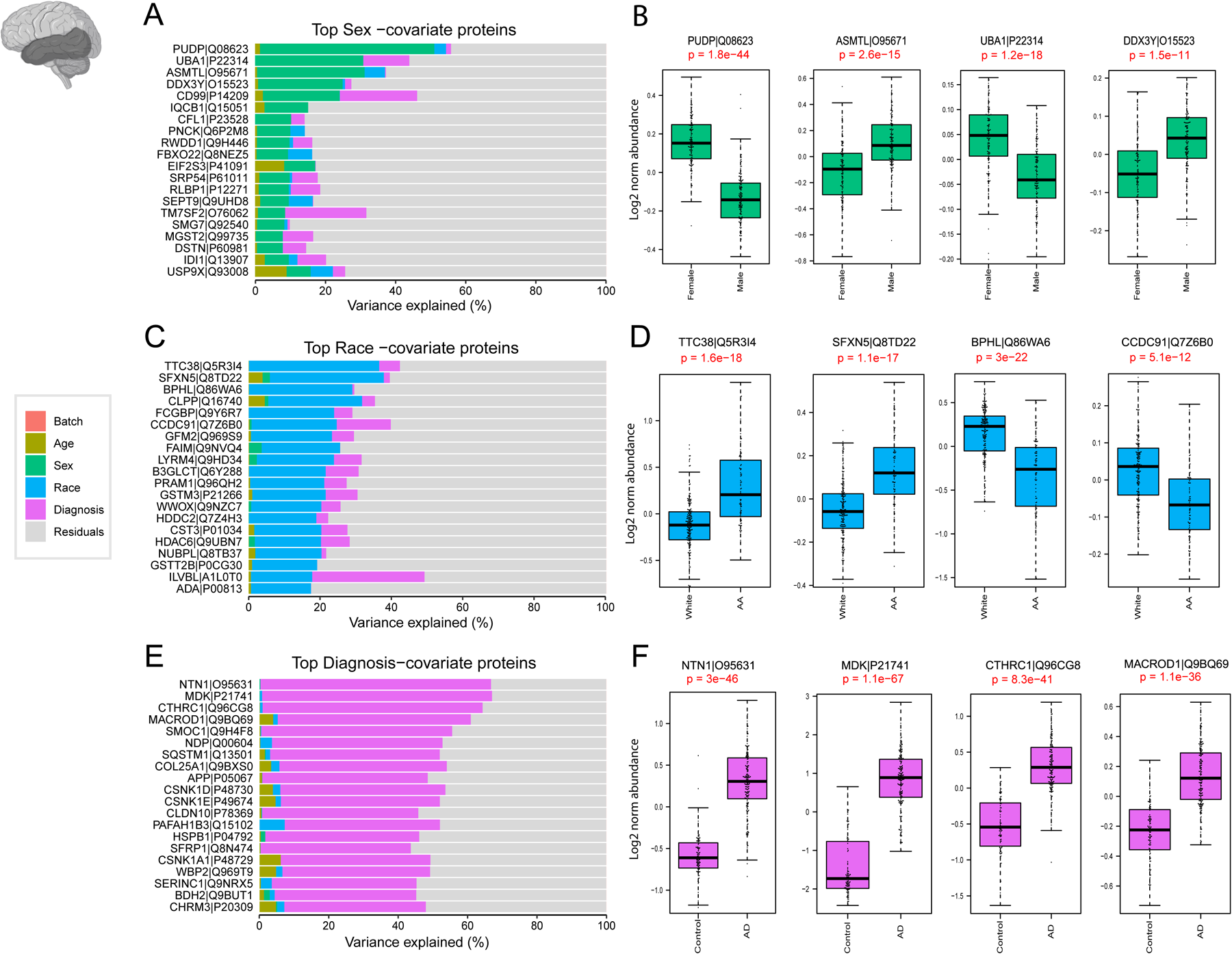
Variance Explained by individual Characteristics in STG Tissues. The bar plots in panels **A, C, and E** illustrate the partitioning of total variance for each protein into fractions attributable to different dimensions of variation in the STG samples. **A**. Top-ranking proteins contributing to sex differences in the dataset were identified through variance partitioning and are presented as bar plots, showing the proportion of variance attributable to sex. Boxplots in panel **B** demonstrate the log2 normal abundance levels of four selected proteins exhibiting significant differences between males and females (p < 0.05). **C.** Bar plots display the top proteins with fraction of total variance attributed to race differences in the STG dataset. Boxplots in panel **D** illustrate the log2 normal abundance levels of four selected proteins demonstrating significant differences between African American individuals and individuals of other races (p < 0.05). **E.** Bar plots identify the top AD-associated proteins with fraction of total variance attributed to AD diagnosis within the STG dataset. Boxplots in Panel F display the log2 normal abundance levels of four selected proteins exhibiting significant differences between AD samples and controls, as well as other diagnostic categories (p < 0.05).

Initially, the relative contribution of sex to the total observed variance was examined. Proteins such as CD99, PUDP, and UBA1, which are associated with the X chromosome and known to be highly abundant in females (36, 37), contributed significantly to the observed variance (**Fig. 4A**, **Fig. 5A**). Conversely, EIF1Y, DDX3Y, and USP9Y, linked to the Y chromosome and known for their high levels in males (36, 37), also played a role in explaining the observed variance. Subsequent analysis confirmed significant differences (P< 0.05) in protein levels between males and females (**Fig. 4B**, **Fig. 5B**), further reinforcing the importance of sex as a determinant of proteomic variability in our dataset.

Key proteins associated with self-reported African American race, such as BPHL, FAIM, GFM2, and CLPP were identified through variance partition analysis in both frontal and temporal cortex samples (**Fig. 4C and Fig. 5C**). In addition, proteins associated with African American race in the temporal cortex displayed a different rank order compared to frontal cortex proteins (**Fig. 5C**). Further examination highlighted significantly higher levels of proteins like FAIM and CLPP in African American individuals. In contrast, the protein KCTD21 exhibited markedly lower levels within this demographic. (**Fig. 4D and Fig. 5D**).

A parallel analysis explored proteins contributing to differences in AD diagnosis within the frontal and temporal cortex. Consistent with existing literature (22, 38, 39), top-ranking proteins associated with AD, included amyloid precursor protein (APP), which has been shown to correlate with β-amyloid plaques burden in brain (22), as well as other amyloid-associated matrisome proteins, CTHRC1, SMOC1, MDK, and NTN1, that all exhibited significantly higher levels among AD cases (40, 41)(**Fig. 4E and 5E**). Notably, some proteins regionally unique to AD diagnosis emerged among the 20 top ranked proteins in the temporal cortex samples including NDP and MACROD1 (**Fig. 5E**).

This comprehensive analysis not only supports the technical validation of our proteomics data but also provides insights into the molecular basis of sex, race, and AD-associated variations in human brain proteomic data.

### 3.3. Correlation between Amyloid and Tau Abundance in Human Brain Proteome and AD Neuropathology

The core pathological hallmarks of AD include β-amyloid (Aβ) plaque and hyperphosphorylated tau neurofibrillary tangle accumulation in the brain (42, 43). As a quality control measure for our dataset, we assessed the levels of the amyloid precursor proteins (APP) and the microtubule associated protein tau (MAPT) using TMT-MS.

The proteomic quantification of APP revealed significantly higher levels in AD cases (p < 0.05) (**Fig. 6A**). Although amyloid precursor protein (APP) is typically a full-length transmembrane protein consisting of 695 amino acid residues in the brain, numerous studies have demonstrated that its levels correlate with Aβ plaque burden in the brain, likely due to the abundance of the β-amyloid peptides in AD brains (44, 45). As part of the amyloid cascade APP undergoes proteolytic cleavage to form Aβ species, which further aggregates to form the characteristic plaques observed in AD brains (46, 47). A further analysis was conducted to measure the correlation between proteomics levels of APP in the frontal cortex and Consortium to Establish a Registry for Alzheimer’s Disease (CERAD) score. The CERAD scores, ranging from 0 to 3, define the extent of neuritic plaques and diffuse plaques in brain tissue, providing a measure of amyloid pathology (48). A total of 980 DLPFC samples from unique individuals were analyzed Our analysis unveiled a stepwise increase in proteomics measurements of APP as CERAD score increased (**Fig. 6B**). This signifies a strong association between CERAD scoring and proteomic quantification of APP levels.

**Fig 6.**
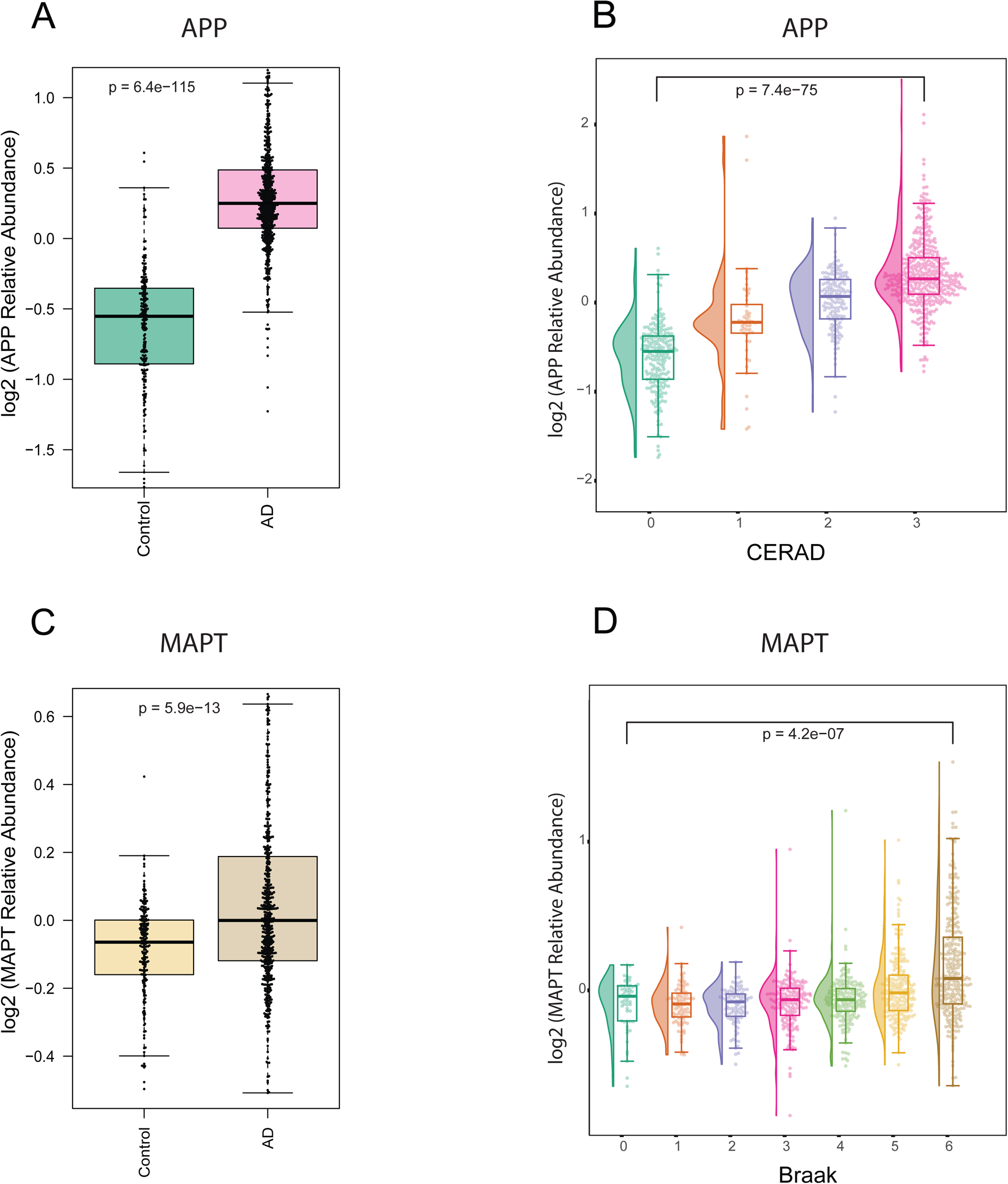
Correlation between proteomic Tau and APP measurements with Braak and CERAD pathological scoring. **A.** Box plots depicting the relative abundance of APP across AD (pink) and control (green) in DLPFC tissue samples (adjusted ANOVA p-value < 0.05). **B.** Raincloud plots depict group differences in the relative abundance of Amyloid Precursor Protein (APP) (Y-axis) across distinct CERAD stages (X-axis) in DLPFC tissues. The analysis revealed a stepwise increase in the median APP levels with ascending CERAD classifications, indicating a progressive trend in APP abundance corresponding to different CERAD groups (score 1: green, score 2: orange, score 3: purple, score 4: pink). **C.** Box plots depicting the relative abundance of MAPT across AD (brown) and control (yellow) in DLPFC tissue samples (adjusted ANOVA p-value < 0.05). **D.** Raincloud plots illustrate the group differences in the relative abundance of Microtubule-associated protein tau (MAPT) (Y-axis) across distinct Braak stages (X-axis) in DLPFC tissues. The Braak stages range from 0 to 6, with corresponding colors representing different stages (0: dark green, 1: orange, 2: purple, 3: pink, 4: light green, 5: yellow, 6: brown). Notably, the analysis highlights elevated MAPT levels at Braak stages 5 and 6, aligning with the expected increase in tau tangles in later stages of Braak in the frontal cortex.

Our proteomic analysis of MAPT, the precursor of tau protein, demonstrated significantly elevated levels in AD cases (P < 0.05) (**Fig. 6C**) likely driven by peptides derived from the Microtubule Binding Region (MTBR), a critical domain associated with tau aggregation and the formation of neurofibrillary tangles (49, 50). A similar approach was conducted to assess the association between the levels of measured MAPT in the frontal cortex and Braak staging. Braak staging ranges from 0 to 6, reflecting the extent of neurofibrillary tangle pathology in different brain regions, with higher scores indicating more advanced stages of AD pathology (51). Regionally, staging starts with the entorhinal cortex and progresses through various regions of the brain, culminating in the neocortex. Proteomic measures of MAPT levels among 980 unique individuals exhibited higher levels mainly in advanced Braak stages. This observation underscores the linkage between proteomic measures of MAPT and tau pathology progression (**Fig. 6D**). In addition, the association between MAPT levels and Braak staging may be influenced by regional differences in tau pathology. Specifically, neurofibrillary tangles are predominantly encountered in the neocortex in higher Braak stages. Therefore, the observed elevation in MAPT levels in individuals with advanced Braak stages could be attributed to the assessment of tau levels in neocortical samples, where tau tangle pathology is pronounced.

Our findings provide valuable insights into the concordance between proteomics measurements and established AD pathology scoring systems.

### 3.4. Association between *APOE4* Genotype and APOE4 Protein Abundance in Human Brain Proteome

In addition to Aβ plaque and tau tangles, the pathophysiology of AD is closely linked with *APOE* genotype (52, 53). *APOE* features three genetic variants (ε2, ε3, ε4), each associated with varying degrees of AD risk, amongst all the *APOE* ε4 allele represents a major genetic risk factor for non-dominantly inherited AD (53, 54). APOE protein variants can be differentiated by arginine-to-cysteine changes that can be detected and measured at the peptide level (55). As part of data validation, we measured the abundance of an APOE4 specific tryptic peptide (LGADMEDVR) and associated it with genotype data for *APOE* across 920 unique individuals where genotyping information was available in DLPFC samples. A similar analysis was employed on 244 unique individuals in STG samples. As expected, given the unique change in protein sequence, the mean fold change of APOE4 protein abundance between *APOE* ε*4* carriers and non-carriers was > 8-fold (**Fig. 7A and 7C**). While it was expected that the APOE4 peptide signal would not be present in non-*APOE* ε4 carriers, cases without an ε4 allele likely still exhibited residual signals for the protein abundance. These signals could be attributed to background chemical noise, possibly stemming from TMT isotope impurity. However, there were a limited number of false positive (non-*APOE* ε4 carriers with APOE4 signal) proteotypes (< 2.0%) observed between expected *APOE* genotype and APOE4 peptide levels, mainly in DLPFC samples. Specifically, we observed 14 number of individuals among DLPFC samples and 2 individuals among STG samples with *APOE* ε3 genotype that had levels of APOE4 equivalent to individuals with *APOE* ε4 genotypes. This could also be due potentially some anticipated errors genotyping. (**Fig. 7B and 7D**). These samples could be removed from further analysis as appropriate. Nevertheless, approximately 97-98% of samples appeared to have the correct *APOE* ε*4* genotype based on APOE4 “proteotype”.

**Fig 7.**
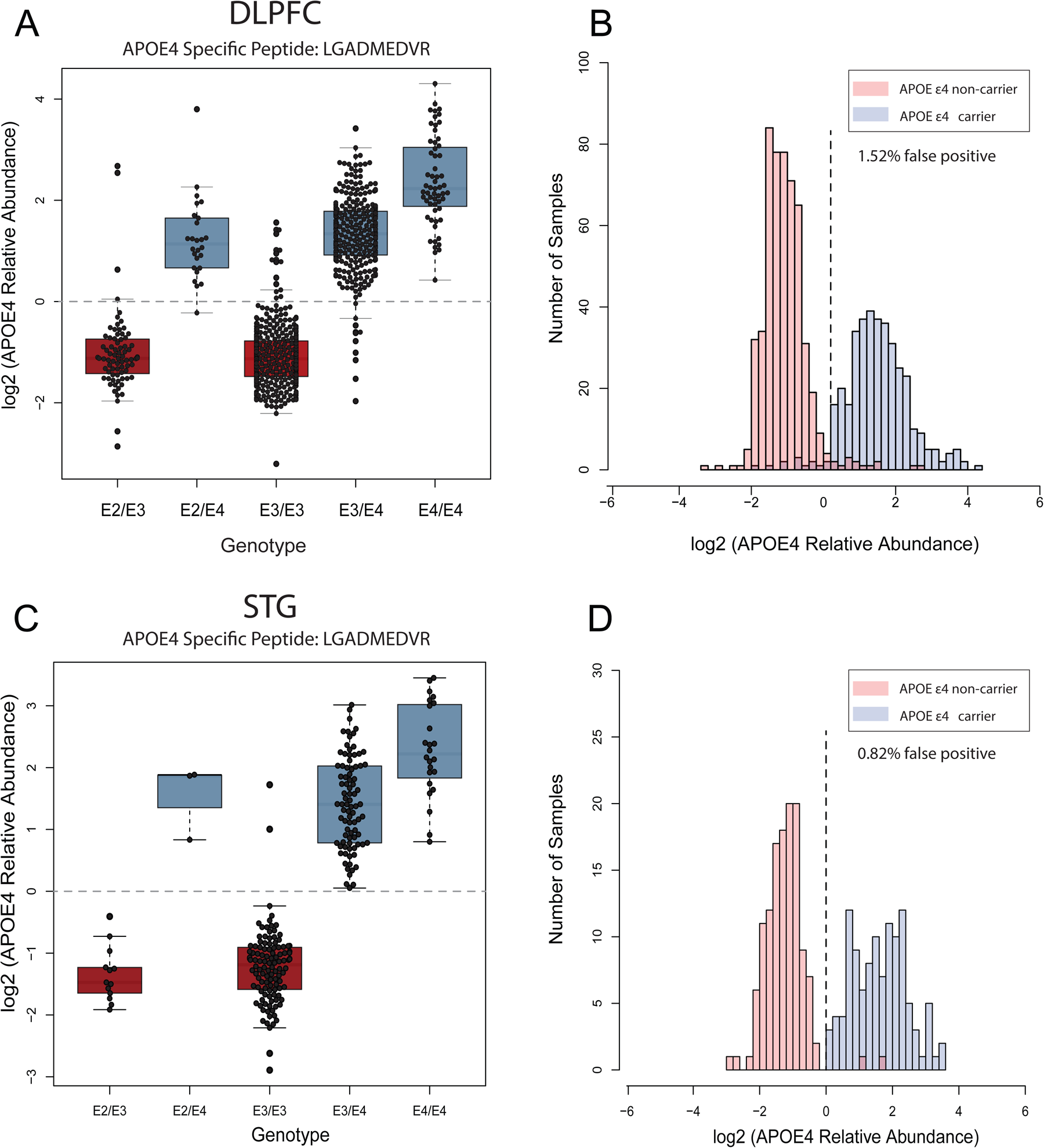
The association between APOE4 genotype and prototype across DLPFC and STG samples. **A.** The boxplots of log2 normal abundance of APOE4 protein measured by TMT-MS across each APOE genotype reveal a high APOE4 abundance among APOE ε4 carriers among 920 unique DLPFC tissue samples. **B.** Histogram of APOE4 log2 normal abundance among DLPFC samples (Y-axis) across ε4 allele presence (red) and non-presence (blue) (X-axis). **C**. The boxplots of log2 normal abundance of APOE4 protein measured by TMT-MS across 244 STG unique tissue samples reveal a high APOE4 abundance among APOE ε4 carriers. **D.** Histogram of APOE4 log2 normal abundance among STG samples (Y-axis) across ε4 allele presence (red) and non-presence (blue) (X-axis). high levels of APOE4 abundance were observed in cases with the ε4 allele combination in both cortices, a few discrepancies between APOE4 genotyping and prototyping (purple) were depicted. These inconsistencies may be attributed to various factors, including mis-genotyping or potential technical challenges in mass spectrometry measurements, such as isotope impurity and low signal-to-noise ratio in specific samples.

## 4. Discussion

Here, we present a comprehensive large-scale deep proteome analysis using TMT-MS on 1105 DLPFC and 280 STG brain tissues. This dataset covered approximately 10,000 proteins from a racially and ethnically diverse cohort comprised of AD and controlled aging brain tissues. In addition, quality control measures were implemented to ensure the validity of our dataset for subsequent analysis. Consistent with the literature (22, 36, 37, 40, 41), our analysis identified top proteins associated with sex, race, and AD diagnosis. Additionally, quantified levels of MAPT and APP showed strong associations with neuropathology scores of Braak and CERAD, respectively. Moreover, the protein abundance of APOE4 was consistent with *APOE* genotyping of the measured samples. These analyses underscore the validity of our data and quality control measures with respect to independent measures of AD pathology and genotype.

This study serves as the data resource for the brain proteome that is also complemented by paired genomics and RNA-seq analyses on these same cases, as supported by the AMP-AD diversity initiative (17, 25). Moving forward, our study sets the stage for future investigations aimed at addressing existing knowledge gaps and advancing our understanding of AD across, age, race, sex and ethnicity. Below we describe several use cases in which this proteomic dataset can be used to address these gaps.

### 4.1. Differential abundance by diagnosis, sex, race and APOE4 genotype

Race, sex, and APOE genotype have been shown to impact AD pathologies, including tau and amyloid in both brain tissue and biofluids (13, 56–61). This large-scale and racially diverse brain tissue proteome dataset provides a valuable resource for investigating differential protein abundance associated with AD diagnosis, sex, race, and APOE and their interactions across brain regions. Researchers can leverage this dataset to identify proteins that exhibit significant differences in abundance between AD cases and controls, as well as between different sexes and racial/ethnic groups. Understanding these differences in proteins influenced by these underlying traits may offer insights into the molecular mechanisms underlying AD risk and pathological burden. Moreover, exploring race-specific protein modulations can contribute to a more comprehensive understanding of disease pathogenesis and aid in the development of targeted interventions tailored to diverse populations.

### 4.2. Network analysis

Unbiased proteomics of the human brain in AD, coupled with network analysis is a valuable approach for organizing and reducing large-scale complex, protein expression matrices data into groups or “modules” of proteins that highly correlate across tissues (28, 30, 62). We and others have shown that these modules reflect various biological functions with cell-type specificity linked to AD pathology (22, 23, 30, 63). Using this approach modules could reveal potential associations between sex, race, APOE genotype and AD diagnosis, shedding light on intersecting biological processes that contribute to disease susceptibility. Furthermore, bulk RNA-seq analysis will be available on majority of these same tissues profiles by proteomics, which will allow for integrated network analyses to compare transcript expression to protein level abundance which are not generally well correlated in human brain tissues (22, 24).

### 4.3. Mapping post-translational modifications

The phosphorylation of tau and proteolytic cleavage of APP into Aβ species are pathological hallmarks of AD and have important roles in disease progression and pathogenesis (42, 43). Other PTMs have also been described as altering the brain proteome in AD (64, 65). Although we did not specifically enrich with antibodies or chemical approaches like immobilized metal affinity chromatography (IMAC) for phosphorylated peptides, the raw MS data can be re-searched to determine if high abundance PTMs like phosphorylation on tau or cleavage of APP to amyloid species (Aβ40, 42 or other species) are altered in these tissues across race.

### 4.4. Proteogenomics

Majority of these tissues profiled by proteomics in this study will have paired whole genome sequencing (WGS). Notably, this offers an opportunity to investigate protein quantitative trait loci or pQTLs to estimate effects of genetic variants on protein abundance (66). Furthermore, integrating AD genome-wide association studies (GWAS) with these pQTLs can be used to pinpoint causal genes that confer AD risk through their effects on brain protein abundance. This approach is referred to as proteome-wide association studies (PWAS), which can now be done with African American GWAS summary statistics AD or related dementia (67, 68). Paired RNA-seq and proteomics from these same tissues can also be used to identify splicing defects in AD that generate alternative protein isoforms occurring in the brain across different disease states and ancestries (69, 70). Understanding how alternative splicing contributes to AD pathophysiology and its intersection with demographic factors may uncover novel disease mechanisms and identify splice variants as potential biomarkers or therapeutic targets.

### 4.5. Limitations and Future Directions

While our study provides valuable insights into the proteomic landscape of AD, several caveats and limitations should be considered. First, it is essential to acknowledge that proteomics data represent a snapshot of protein abundance at a particular point in time and may not capture dynamic changes in protein expression over the course of disease progression. Additionally, although efforts were made to minimize technical variability through rigorous quality control measures, the inherent complexity of brain tissues and potential confounding factors such as comorbidities may introduce biases or artifacts into the dataset. In addition, a few discrepancies were noted, for example, the number of controls in our study was not matched with the number of AD cases, resulting in fewer control cases. Moreover, the lack of post-mortem interval (PMI) information for all samples is another limitation of our study. Without PMI data, we were unable to account for the potential effects of PMI on protein degradation. It is also important to mention that the interpretation of race-specific protein modulations should be approached with caution, as the biological basis underlying these differences remains incompletely understood. Further validation studies and replication in independent cohorts are warranted to confirm and extend our findings. Future proteomic studies on biofluids from diverse participant from CSF and plasma will be warranted to understand how these changes in the post-mortem brain are reflected in the periphery and are prognostic for AD. Ultimately, integrated multi-omic datasets across tissues and biofluids, will be needed for further investigation into how AD heterogeneity varies across different ethnoracial backgrounds.

## Data Availability

Raw data files and clinical metadata are available at https://doi.org/10.7303/syn53420674. Search results and database, sample-to-TMT channel information, normalized data are available at https://doi.org/10.7303/syn55225561. Summary data is also available at the ShinyApp https://telomere.biochem.emory.edu/diversity/.

## Acknowledgement

We would like to thank all the participants, brain donors, and their families, without whom this study was impossible. The results published here are based on data available in the AD Knowledge Portal (https://adknowledgeportal.org). The Mayo RNAseq study data was led by Dr. Nilüfer Ertekin-Taner, Mayo Clinic, Jacksonville, FL, as part of the multi-PI U01 AG046139 (MPIs Golde, Ertekin-Taner, Younkin, Price) using samples from The Mayo Clinic Brain Bank. Data collection was supported through funding by NIA grants P50 AG016574, R01 AG032990, U01 AG046139, R01 AG018023, U01 AG006576, U01 AG006786, R01 AG025711, R01 AG017216, R01 AG003949, P30AG072979, P01AG066597, U19AG062418, U01AG061357, RF1AG062181, P30AG066511 CurePSP Foundation, and support from Mayo Foundation. Study data included samples collected through the Sun Health Research Institute Brain and Body Donation Program of Sun City, Arizona, USA. The Brain and Body Donation Program has been supported by the National Institute of Neurological Disorders and Stroke (U24 NS072026 National Brain and Tissue Resource for Parkinson’s Disease and Related Disorders), the National Institute on Aging (P30 AG019610 and P30AG072980, Arizona Alzheimer’s Disease Center), the Arizona Department of Health Services (contract 211002, Arizona Alzheimer’s Research Center), the Arizona Biomedical Research Commission (contracts 4001, 0011, 05-901 and 1001 to the Arizona Parkinson’s Disease Consortium) and the Michael J. Fox Foundation for Parkinson’s Research. We would like to thank John Q. Trojanowski (deceased) for his leadership at the Center for Neurodegenerative Disease Research, which helped make acquiring samples from University of Pennsylvania Brain Bank possible.

Additional support for these studies was provided by the NINDS grant R01-NS080820 (NET), NIA grant R01-AG061796 (NET), NIA grant U19-AG074879 (NET), Alzheimer’s Association Zenith Fellows Award (NET) and P30AG072979 (EBL), P01AG066597 (EBL), U19AG062418 (EBL). We thank the Mayo Clinic Genome Analysis Core (GAC), Co-Directors Julie M. Cunningham, PhD and Eric Wieben, PhD, and supervisor Julie Lau, for their collaboration in the collection of omics data. The Rush Alzheimer’s Disease Center contributed data and brain biospecimens from the Religious Orders Study and Rush Memory and Aging Project (ROSMAP), the Minority Aging Research Study, and the African American and Latino Cores with funding from P30AG10161, P30AG72975, R01AG15819, R01AG17917, R01AG22018, U01AG46152, and U01AG61356.

## Notes

https://doi.org/10.7303/syn55225561

https://telomere.biochem.emory.edu/diversity/

https://doi.org/10.7303/syn53420674

### Competing Interest Statement

The authors have declared no competing interest.

## Bibliography

1. 1. Estimation of the global prevalence of dementia in 2019 and forecasted prevalence in 2050: an analysis for the Global Burden of Disease Study 2019. The Lancet Public health. 2022;7(2):e105–e25.

2. 2023 Alzheimer’s disease facts and figures. Alzheimer’s & dementia : the journal of the Alzheimer’s Association. 2023;19(4):1598–695.

3. Alzheimer’s disease facts and figures. Alzheimer’s & dementia : the journal of the Alzheimer’s Association. 2010;6(2):158–94.

4. Chin AL, Negash S, Hamilton R. Diversity and disparity in dementia: the impact of ethnoracial differences in Alzheimer disease. Alzheimer disease and associated disorders. 2011;25(3):187–95.

5. Le Guen Y, Raulin A-C, Logue MW, Sherva R, Belloy ME, Eger SJ, et al. Association of African Ancestry–Specific APOE Missense Variant R145C With Risk of Alzheimer Disease. JAMA. 2023;329(7):551–60.

6. Lourida I, Hannon E, Littlejohns TJ, Langa KM, Hyppönen E, Kuzma E, et al. Association of Lifestyle and Genetic Risk With Incidence of Dementia. Jama. 2019;322(5):430–7.

7. Almeida RP, Schultz SA, Austin BP, Boots EA, Dowling NM, Gleason CE, et al. Effect of Cognitive Reserve on Age-Related Changes in Cerebrospinal Fluid Biomarkers of Alzheimer Disease. JAMA neurology. 2015;72(6):699–706.

8. Caunca MR, Odden MC, Glymour MM, Elfassy T, Kershaw KN, Sidney S, et al. Association of Racial Residential Segregation Throughout Young Adulthood and Cognitive Performance in Middle-aged Participants in the CARDIA Study. JAMA neurology. 2020;77(8):1000–7.

9. Hunt JFV, Buckingham W, Kim AJ, Oh J, Vogt NM, Jonaitis EM, et al. Association of Neighborhood-Level Disadvantage With Cerebral and Hippocampal Volume. JAMA neurology. 2020;77(4):451–60.

10. Lamar M, Lerner AJ, James BD, Yu L, Glover CM, Wilson RS, et al. Relationship of Early-Life Residence and Educational Experience to Level and Change in Cognitive Functioning: Results of the Minority Aging Research Study. The journals of gerontology Series B, Psychological sciences and social sciences. 2020;75(7):e81–e92.

11. National Research Council Panel on Race E, Health in Later L. The National Academies Collection: Reports funded by National Institutes of Health. In: Anderson NB, Bulatao RA, Cohen B, editors. Critical Perspectives on Racial and Ethnic Differences in Health in Late Life. Washington (DC): National Academies Press (US) Copyright © 2004, National Academy of Sciences.; 2004.

12. Sisco S, Gross AL, Shih RA, Sachs BC, Glymour MM, Bangen KJ, et al. The role of early-life educational quality and literacy in explaining racial disparities in cognition in late life. The journals of gerontology Series B, Psychological sciences and social sciences. 2015;70(4):557–67.

13. Modeste ES, Ping L, Watson CM, Duong DM, Dammer EB, Johnson ECB, et al. Quantitative proteomics of cerebrospinal fluid from African Americans and Caucasians reveals shared and divergent changes in Alzheimer’s disease. Molecular neurodegeneration. 2023;18(1):48.

14. Morris JC, Schindler SE, McCue LM, Moulder KL, Benzinger TLS, Cruchaga C, et al. Assessment of Racial Disparities in Biomarkers for Alzheimer Disease. JAMA neurology. 2019;76(3):264–73.

15. Garrett SL, McDaniel D, Obideen M, Trammell AR, Shaw LM, Goldstein FC, et al. Racial Disparity in Cerebrospinal Fluid Amyloid and Tau Biomarkers and Associated Cutoffs for Mild Cognitive Impairment. JAMA network open. 2019;2(12):e1917363.

16. Howell JC, Watts KD, Parker MW, Wu J, Kollhoff A, Wingo TS, et al. Race modifies the relationship between cognition and Alzheimer’s disease cerebrospinal fluid biomarkers. Alzheimer’s research & therapy. 2017;9(1):88.

17. Hodes RJ, Buckholtz N. Accelerating Medicines Partnership: Alzheimer’s Disease (AMP-AD) Knowledge Portal Aids Alzheimer’s Drug Discovery through Open Data Sharing. Expert opinion on therapeutic targets. 2016;20(4):389–91.

18. Bayer TA. Proteinopathies, a core concept for understanding and ultimately treating degenerative disorders? European neuropsychopharmacology : the journal of the European College of Neuropsychopharmacology. 2015;25(5):713–24.

19. Bai B, Vanderwall D, Li Y, Wang X, Poudel S, Wang H, et al. Proteomic landscape of Alzheimer’s Disease: novel insights into pathogenesis and biomarker discovery. Molecular neurodegeneration. 2021;16(1):55.

20. Macklin A, Khan S, Kislinger T. Recent advances in mass spectrometry based clinical proteomics: applications to cancer research. Clinical Proteomics. 2020;17(1):17.

21. Rayaprolu S, Higginbotham L, Bagchi P, Watson CM, Zhang T, Levey AI, et al. Systems-based proteomics to resolve the biology of Alzheimer’s disease beyond amyloid and tau. Neuropsychopharmacology : official publication of the American College of Neuropsychopharmacology. 2021;46(1):98–115.

22. Johnson ECB, Carter EK, Dammer EB, Duong DM, Gerasimov ES, Liu Y, et al. Large-scale deep multi-layer analysis of Alzheimer’s disease brain reveals strong proteomic disease-related changes not observed at the RNA level. Nat Neurosci. 2022;25(2):213–25.

23. Seyfried NT, Dammer EB, Swarup V, Nandakumar D, Duong DM, Yin L, et al. A Multi-network Approach Identifies Protein-Specific Co-expression in Asymptomatic and Symptomatic Alzheimer’s Disease. Cell Systems. 2017;4(1):60–72.e4.

24. Tasaki S, Xu J, Avey DR, Johnson L, Petyuk VA, Dawe RJ, et al. Inferring protein expression changes from mRNA in Alzheimer’s dementia using deep neural networks. Nature Communications. 2022;13(1):655.

25. Reddy JS, Heath L, Linden AV, Allen M, Lopes KdP, Seifar F, et al. Bridging the Gap: Multi-Omics Profiling of Brain Tissue in Alzheimer’s Disease and Older Controls in Multi-Ethnic Populations. bioRxiv. 2024.

26. Guo Q, Ping L, Dammer EB, Duong DM, Yin L, Xu K, et al. Global analysis of the heparin-enriched plasma proteome captures matrisome-associated proteins in Alzheimer’s disease. bioRxiv. 2023.

27. Marks JD, Ayuso VE, Carlomagno Y, Yue M, Todd TW, Hao Y, et al. TMEM106B core deposition associates with TDP-43 pathology and is increased in risk SNP carriers for frontotemporal dementia. Science translational medicine. 2024;16(730):eadf9735.

28. Wojtas AM, Dammer EB, Guo Q, Ping L, Shantaraman A, Duong DM, et al. Proteomic Changes in the Human Cerebrovasculature in Alzheimer’s Disease and Related Tauopathies Linked to Peripheral Biomarkers in Plasma and Cerebrospinal Fluid. medRxiv : the preprint server for health sciences. 2024.

29. Käll L, Canterbury JD, Weston J, Noble WS, MacCoss MJ. Semi-supervised learning for peptide identification from shotgun proteomics datasets. Nature methods. 2007;4(11):923–5.

30. Johnson ECB, Dammer EB, Duong DM, Ping L, Zhou M, Yin L, et al. Large-scale proteomic analysis of Alzheimer’s disease brain and cerebrospinal fluid reveals early changes in energy metabolism associated with microglia and astrocyte activation. Nat Med. 2020;26(5):769–80.

31. Robins C, Liu Y, Fan W, Duong DM, Meigs J, Harerimana NV, et al. Genetic control of the human brain proteome. American journal of human genetics. 2021;108(3):400–10.

32. Wingo AP, Fan W, Duong DM, Gerasimov ES, Dammer EB, Liu Y, et al. Shared proteomic effects of cerebral atherosclerosis and Alzheimer’s disease on the human brain. Nature neuroscience. 2020;23(6):696–700.

33. Dammer EB, Seyfried NT, Johnson ECB. Batch Correction and Harmonization of -Omics Datasets with a Tunable Median Polish of Ratio. Frontiers in systems biology. 2023;3.

34. Yu Y, Zhang N, Mai Y, Ren L, Chen Q, Cao Z, et al. Correcting batch effects in large-scale multiomics studies using a reference-material-based ratio method. Genome biology. 2023;24(1):201.

35. Hout MC, Papesh MH, Goldinger SD. Multidimensional scaling. Wiley interdisciplinary reviews Cognitive science. 2013;4(1):93–103.

36. Armoskus C, Moreira D, Bollinger K, Jimenez O, Taniguchi S, Tsai HW. Identification of sexually dimorphic genes in the neonatal mouse cortex and hippocampus. Brain research. 2014;1562:23–38.

37. Braun AE, Mitchel OR, Gonzalez TL, Sun T, Flowers AE, Pisarska MD, et al. Sex at the interface: the origin and impact of sex differences in the developing human placenta. Biology of sex differences. 2022;13(1):50.

38. Johnson ECB, Bian S, Haque RU, Carter EK, Watson CM, Gordon BA, et al. Cerebrospinal fluid proteomics define the natural history of autosomal dominant Alzheimer’s disease. Nature Medicine. 2023;29(8):1979–88.

39. Levites Y, Dammer EB, Ran Y, Tsering W, Duong D, Abreha M, et al. Aβ Amyloid Scaffolds the Accumulation of Matrisome and Additional Proteins in Alzheimer’s Disease, bioRxiv. 2023.

40. Askenazi M, Kavanagh T, Pires G, Ueberheide B, Wisniewski T, Drummond E. Compilation of reported protein changes in the brain in Alzheimer’s disease. Nature Communications. 2023;14(1):4466.

41. Drummond E, Nayak S, Faustin A, Pires G, Hickman RA, Askenazi M, et al. Proteomic differences in amyloid plaques in rapidly progressive and sporadic Alzheimer’s disease. Acta neuropathologica. 2017;133(6):933–54.

42. Knopman DS, Amieva H, Petersen RC, Chételat G, Holtzman DM, Hyman BT, et al. Alzheimer disease. Nature Reviews Disease Primers. 2021;7(1):33.

43. Kocahan S, Doğan Z. Mechanisms of Alzheimer’s Disease Pathogenesis and Prevention: The Brain, Neural Pathology, N-methyl-D-aspartate Receptors, Tau Protein and Other Risk Factors. Clinical psychopharmacology and neuroscience : the official scientific journal of the Korean College of Neuropsychopharmacology. 2017;15(1):1–8.

44. Bloom GS. Amyloid-β and tau: the trigger and bullet in Alzheimer disease pathogenesis. JAMA neurology. 2014;71(4):505–8.

45. Ittner LM, Götz J. Amyloid-β and tau--a toxic pas de deux in Alzheimer’s disease. Nature reviews Neuroscience. 2011;12(2):65–72.

46. Chen G-f, Xu T-h, Yan Y, Zhou Y-r, Jiang Y, Melcher K, et al. Amyloid beta: structure, biology and structure-based therapeutic development. Acta Pharmacologica Sinica. 2017;38(9):1205–35.

47. Chow VW, Mattson MP, Wong PC, Gleichmann M. An overview of APP processing enzymes and products. Neuromolecular medicine. 2010;12(1):1–12.

48. Mirra SS, Heyman A, McKeel D, Sumi SM, Crain BJ, Brownlee LM, et al. The Consortium to Establish a Registry for Alzheimer’s Disease (CERAD). Part II. Standardization of the neuropathologic assessment of Alzheimer’s disease. Neurology. 1991;41(4):479–86.

49. Dai J, Johnson ECB, Dammer EB, Duong DM, Gearing M, Lah JJ, et al. Effects of APOE Genotype on Brain Proteomic Network and Cell Type Changes in Alzheimer’s Disease. Frontiers in molecular neuroscience. 2018;11:454.

50. Toombs J, Zetterberg H. Untangling the tau microtubule-binding region. Brain : a journal of neurology. 2021;144(2):359–62.

51. Braak H, Braak E. Neuropathological stageing of Alzheimer-related changes. Acta neuropathologica. 1991;82(4):239–59.

52. Blanchard JW, Akay LA, Davila-Velderrain J, von Maydell D, Mathys H, Davidson SM, et al. APOE4 impairs myelination via cholesterol dysregulation in oligodendrocytes. Nature. 2022;611(7937):769-79.

53. Raulin A-C, Doss SV, Trottier ZA, Ikezu TC, Bu G, Liu C-C. ApoE in Alzheimer’s disease: pathophysiology and therapeutic strategies. Molecular neurodegeneration. 2022;17(1):72.

54. de Rojas I, Moreno-Grau S, Tesi N, Grenier-Boley B, Andrade V, Jansen IE, et al. Common variants in Alzheimer’s disease and risk stratification by polygenic risk scores. Nature Communications. 2021;12(1):3417.

55. Watson CM, Dammer EB, Ping L, Duong DM, Modeste E, Carter EK, et al. Quantitative Mass Spectrometry Analysis of Cerebrospinal Fluid Protein Biomarkers in Alzheimer’s Disease. Sci Data. 2023;10(1):261.

56. Desaire H, Stepler KE, Robinson RAS. Exposing the Brain Proteomic Signatures of Alzheimer’s Disease in Diverse Racial Groups: Leveraging Multiple Data Sets and Machine Learning. Journal of proteome research. 2022;21(4):1095–104.

57. Tijms BM, Gobom J, Reus L, Jansen I, Hong S, Dobricic V, et al. Pathophysiological subtypes of Alzheimer’s disease based on cerebrospinal fluid proteomics. Brain : a journal of neurology. 2020;143(12):3776–92.

58. Tijms BM, Gobom J, Teunissen C, Dobricic V, Tsolaki M, Verhey F, et al. CSF Proteomic Alzheimer’s Disease-Predictive Subtypes in Cognitively Intact Amyloid Negative Individuals. Proteomes. 2021;9(3).

59. Tijms BM, Vromen EM, Mjaavatten O, Holstege H, Reus LM, van der Lee S, et al. Cerebrospinal fluid proteomics in patients with Alzheimer’s disease reveals five molecular subtypes with distinct genetic risk profiles. Nature Aging. 2024;4(1):33–47.

60. Walters S, Contreras AG, Eissman JM, Mukherjee S, Lee ML, Choi S-E, et al. Associations of Sex, Race, and Apolipoprotein E Alleles With Multiple Domains of Cognition Among Older Adults. JAMA neurology. 2023;80(9):929–39.

61. Wang YT, Therriault J, Servaes S, Tissot C, Rahmouni N, Macedo AC, et al. Sex-specific modulation of amyloid-β on tau phosphorylation underlies faster tangle accumulation in females. Brain : a journal of neurology. 2023.

62. Langfelder P, Horvath S. WGCNA: an R package for weighted correlation network analysis. BMC bioinformatics. 2008;9:559.

63. Zhang Q, Ma C, Gearing M, Wang PG, Chin L-S, Li L. Integrated proteomics and network analysis identifies protein hubs and network alterations in Alzheimer’s disease. Acta Neuropathologica Communications. 2018;6(1):19.

64. Arakhamia T, Lee CE, Carlomagno Y, Duong DM, Kundinger SR, Wang K, et al. Posttranslational Modifications Mediate the Structural Diversity of Tauopathy Strains. Cell. 2020;180(4):633–44.e12.

65. Bastrup J, Kastaniegaard K, Asuni AA, Volbracht C, Stensballe A. Proteomic and Unbiased Post-Translational Modification Profiling of Amyloid Plaques and Surrounding Tissue in a Transgenic Mouse Model of Alzheimer’s Disease. Journal of Alzheimer’s disease : JAD. 2020;73(1):393–411.

66. Wingo AP, Liu Y, Gerasimov ES, Gockley J, Logsdon BA, Duong DM, et al. Integrating human brain proteomes with genome-wide association data implicates new proteins in Alzheimer’s disease pathogenesis. Nature genetics. 2021;53(2):143–6.

67. Kunkle BW, Schmidt M, Klein H-U, Naj AC, Hamilton-Nelson KL, Larson EB, et al. Novel Alzheimer Disease Risk Loci and Pathways in African American Individuals Using the African Genome Resources Panel: A Meta-analysis. JAMA neurology. 2021;78(1):102–13.

68. Lake J, Warly Solsberg C, Kim JJ, Acosta-Uribe J, Makarious MB, Li Z, et al. Multi-ancestry meta-analysis and fine-mapping in Alzheimer’s disease. Molecular Psychiatry. 2023;28(7):3121–32.

69. Johnson ECB, Dammer EB, Duong DM, Yin L, Thambisetty M, Troncoso JC, et al. Deep proteomic network analysis of Alzheimer’s disease brain reveals alterations in RNA binding proteins and RNA splicing associated with disease. Molecular neurodegeneration. 2018;13(1):52.

70. Raj T, Li YI, Wong G, Humphrey J, Wang M, Ramdhani S, et al. Integrative transcriptome analyses of the aging brain implicate altered splicing in Alzheimer’s disease susceptibility. Nature genetics. 2018;50(11):1584–92.

